## Supplemental Information for "A single pair of pharyngeal neurons functions as a commander to reject high salt in *Drosophila melanogaster*"

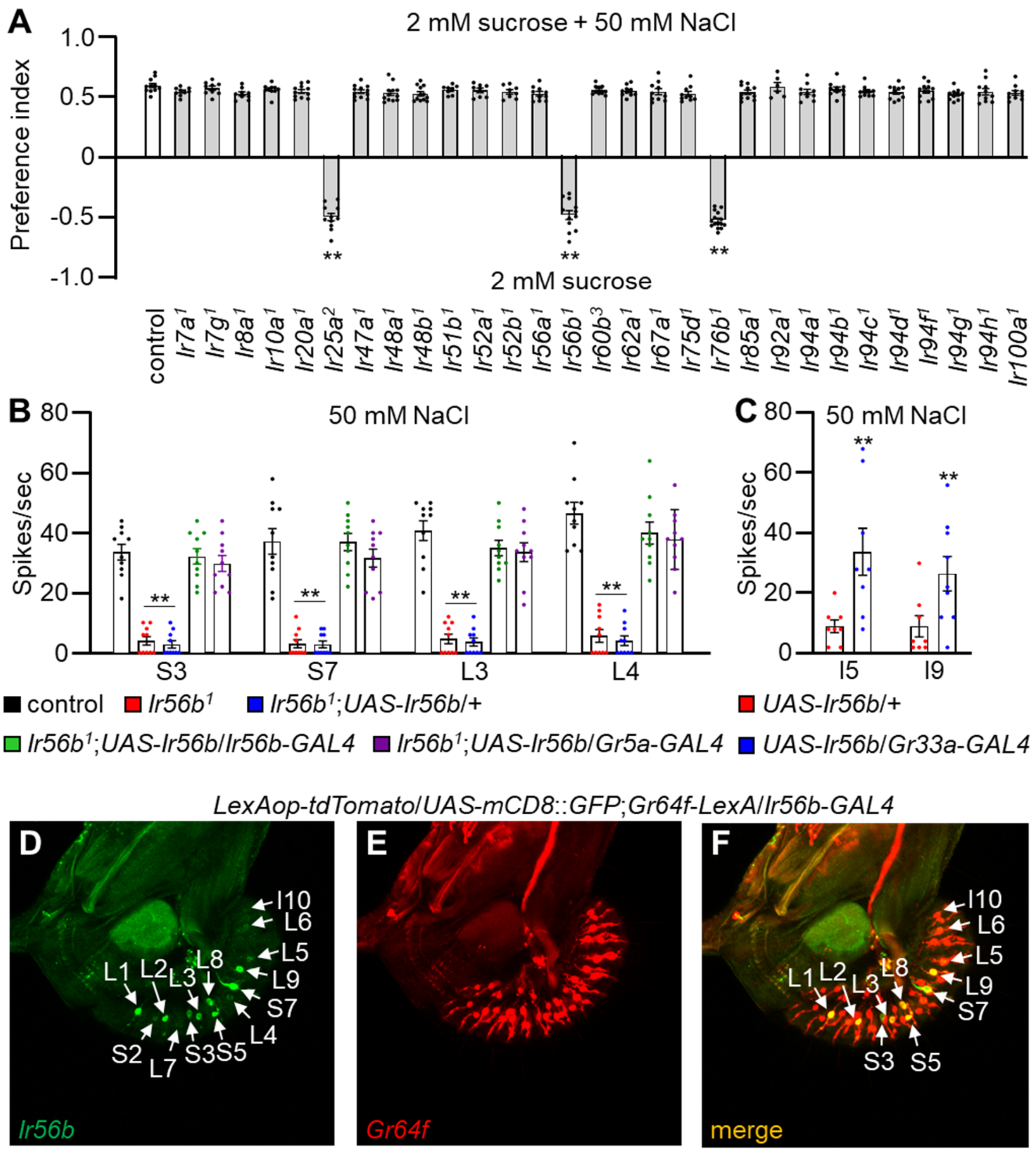


**Figure 1—figure supplement 1.** Requirements for *Ir*s for preferring low salt-containing food. (**A**) Binary food choice assays to assess low salt attraction by comparing *Ir* mutants with the control ($w^{1118}$). n=8–12. (**B**) Tip recordings performed on S3, S7, L3, and L4 sensilla. Shown are comparisons between control, $Ir56b^{1}$, and flies expressing *Ir56b* under control of the *Ir56b-GAL4* or the *Gr5a-GAL4.* n=8–12. (**C**) *UAS-Ir56b* expressed in Class B GRNs using the *Gr33a-GAL4* driver. Tip recordings were conducted on I5 and I9 sensilla of the indicated flies with 50 mM NaCl. n=8. (**D—E**) Immunohistochemistry was performed using anti-GFP and anti-RFP on a labellum from a *LexAop-tdTomato*/*UAS-mCD8*::*GFP*;*Gr64f-LexA*/*Ir56b-GAL4* fly. Multiple sets of data were compared using single-factor ANOVA coupled with Scheffe's *post hoc* test. Statistical significance was relative to the control. Means ± SEMs. **p < 0.01.

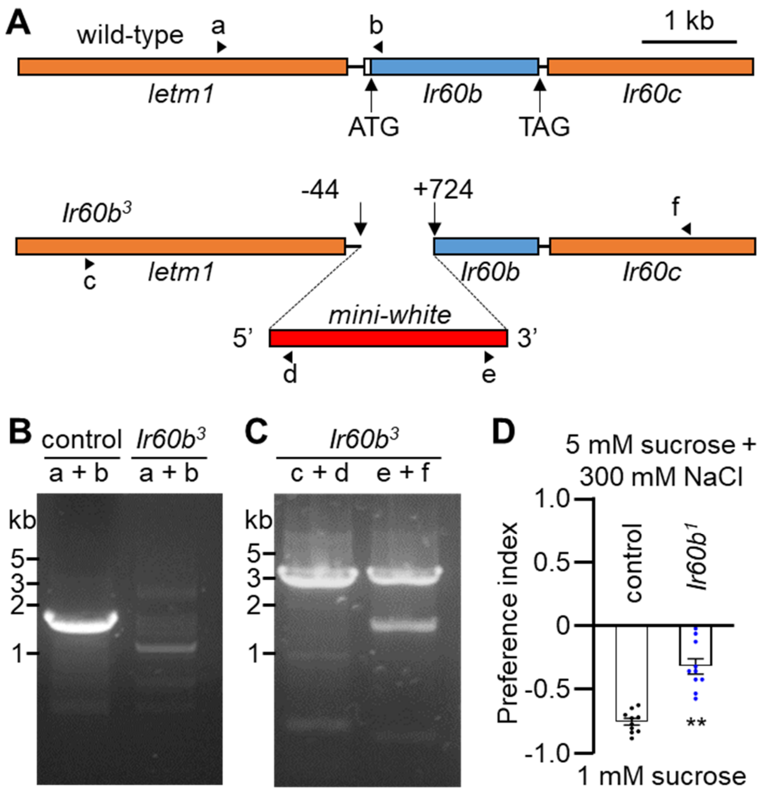


**Figure 1—figure supplement 2.** Gene structure of *Ir60b* locus, generation of *Ir60b³* and behavioral defect of *Ir60b¹* in high salt avoidance. (**A**) Schematic of the *Ir60b* locus and *Ir60b³* allele. The *Ir60b* coding exon is indicated by the blue rectangle. *Ir60b³* was generated by ends-out homologous recombination by removing 768 base pairs as indicated. The red box indicates the insertion of the *mini-white* gene. The arrowheads (a—e) indicate the primers used for the PCR analyses in (**B**) and (**C**). (**B**) Confirmation of the deletion in *Ir60b³* by PCR using primers a and b (primer a: 5'-TTGGTGTTTACTCGAAAACA-3', primer b: 5'-GCATTCAGAATGTATCTTAG-3'). (**C**) Confirmation of the deletion in *Ir60b³* by PCR using primers c and d, and e and f (primer c: 5'-CGAACTGCATGCGCAACAGT-3', primer d: 5'-TTGCTGCCTCCGCGAATTAA-3', primer e: 5'-TGTACTACTCACATTGTTCA-3', primer f: 5'-GATTGTGAGCAGCAGCAGCA-3'). (**D**) Binary food choice assay testing control and *Ir60b¹* flies with 1 mM sucrose versus 5 mM sucrose and 300 mM NaCl. n=10. The pairwise comparison was conducted using a Student *t*-test. Means ± SEMs. **p < 0.01.

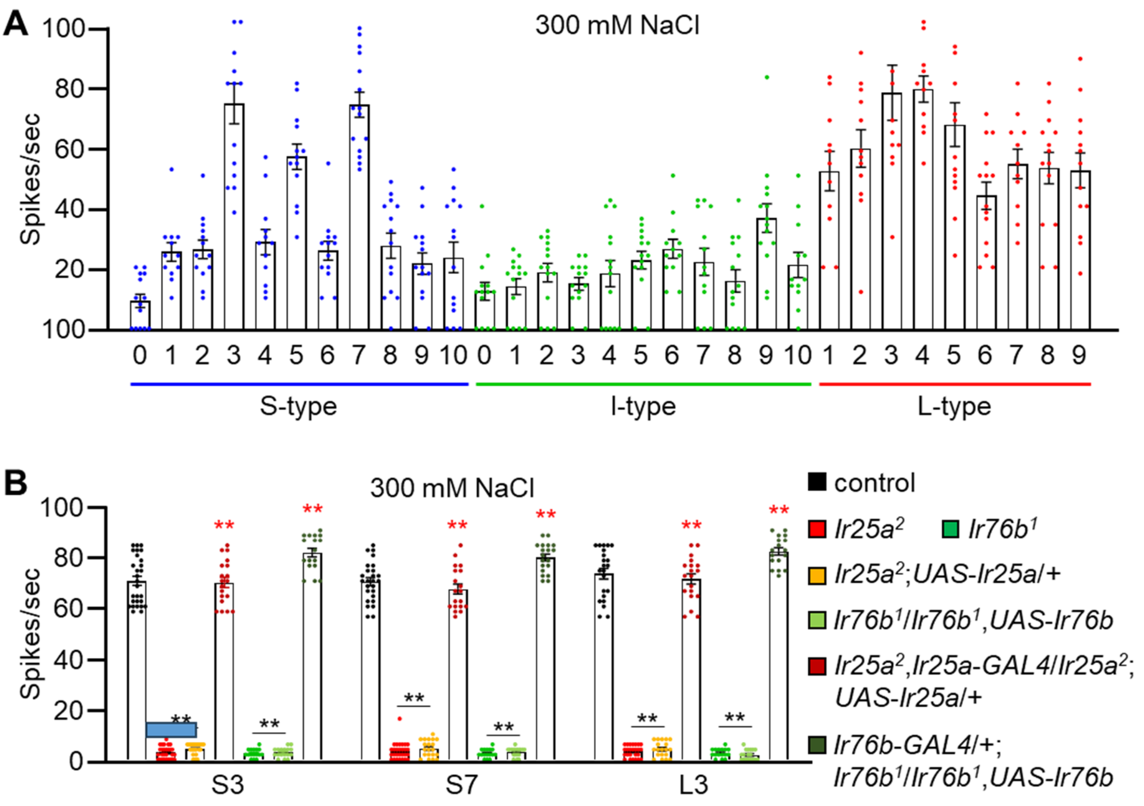


**Figure 2—figure supplement 1.** Assaying action potentials induced by different labellar bristles in response to 300 mM salt using tip recordings. (**A**) Tip recordings were performed on all S-type, I-type and L-type sensilla from control flies. n=12—14. (**B**) Tip recordings conducted on S3, S7, and L3 sensilla from the indicated *Ir25a*$^{2}$ and *Ir76b*$^{1}$ mutants, as well as the mutants expressing wild-type *UAS-Ir25a* and *UAS-Ir76b* transgenes expressed under control of the *Ir25a-GAL4* and the *Ir76b-GAL4*, respectively. n=16–28. Multiple sets of data were compared using single-factor ANOVA coupled with the Scheffe's post hoc test. Statistical significances compared to the control line are indicated by the black asterisks, while the red asterisks indicate significant rescue compared to the corresponding mutant. Means ± SEMs. **p < 0.01.

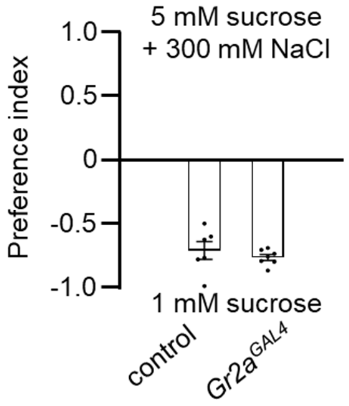


**Figure 2—figure supplement 2.** Two-way solid-food choice assay to assess whether the *Gr2a*$^{GAL4}$ mutant exhibits a deficit in avoidance of high salt. The flies were given a choice between 1 mM sucrose versus 5 mM sucrose plus 300 mM NaCl in alternating wells of microtiter dishes. n=8. The pairwise comparison was conducted using a Student *t*-test. Means ± SEMs.

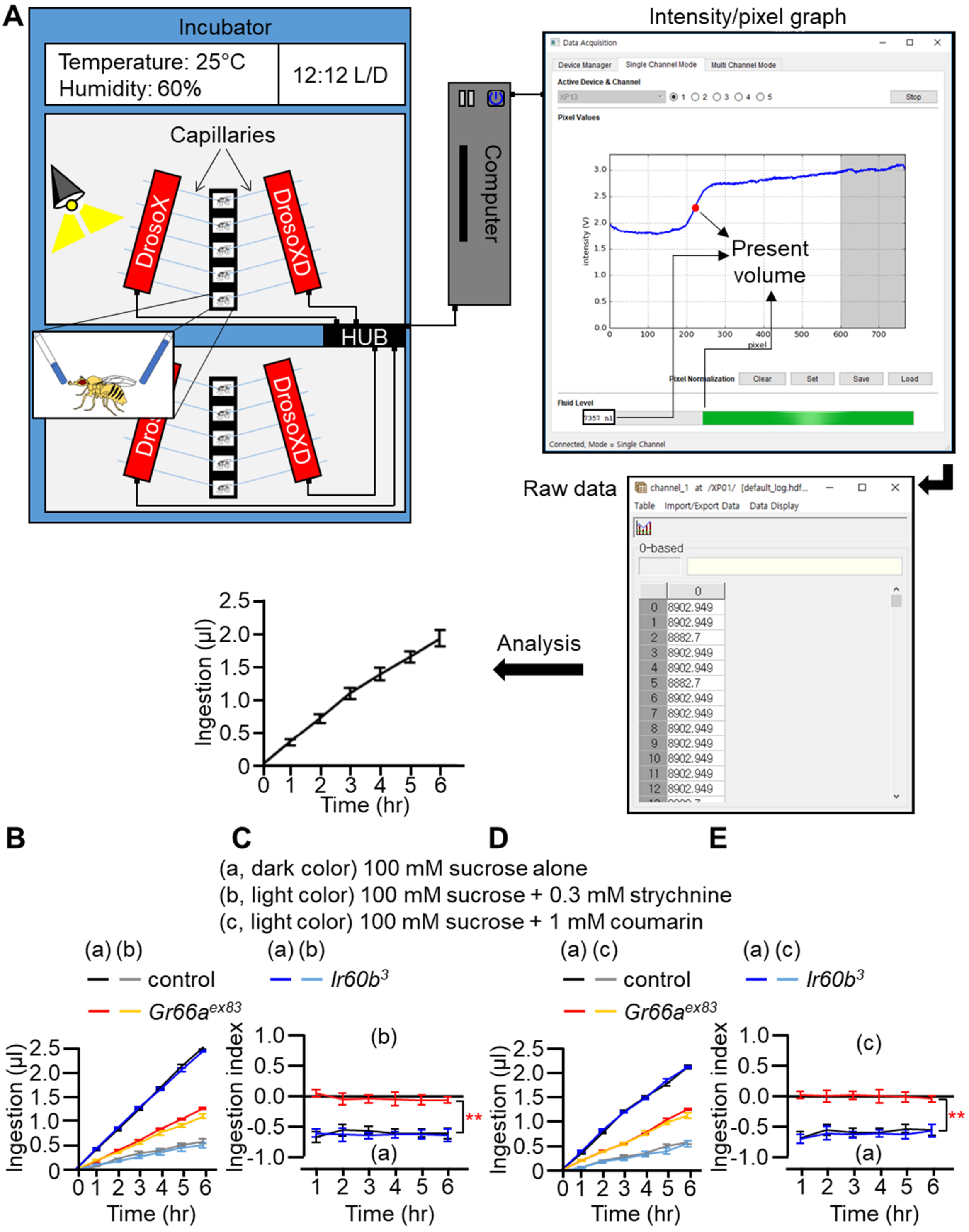


**Figure 3—figure supplement 1.** DrosoX system and measurement of food intake using strychnine and coumarin. (**A**) The DrosoX system is composed of a cassette

designed to accommodate five flies and a monitor that can hold five pairs of capillaries containing different solutions. Flies are exposed to both the DrosoX capillary and DrosoXD capillary options, enabling binary food (solution) choice experiments. A computer records the electrical signal generated by each photodiode, and the light intensity at each pixel in the array was sampled at 8 Hz by a microcontroller on a development board attached to the DrosoX sensor bank. Liquid level readings are acquired at sample rates of 2 Hz. The red dot on the blue line in the intensity/pixel graph indicates the present volume. The raw data is displayed on the monitor with an accuracy of ±5.78 nL. The remaining volumes of solution are recorded based on the difference in optical density between air and the solution during the 6 hr experiment. The formula for calculating ingestion volume is ($Volume_{initial}$ – $Volume_{time\ point}$). (**B—E**) Each fly (control, *Ir60b*$^{3}$, and *Gr66a*$^{ex83}$) was exposed to two capillaries, one of which contained 100 mM sucrose (a), and the other contained 100 mM sucrose and either 0.3 mM strychnine or 1 mM coumarin as indicated (b). (**B** and **D**) Volumes of the two food options consumed by the indicated flies over the course of 6 hrs. (**C** and **E**) Ingestion indexes (I.I) to indicate the relative consumption of the two foods. I.I formula: [Ingestion $volume_{(b)\ or\ (c)}$ – Ingestion $volume_{(a)}$]/ [Ingestion $volume_{(b)\ or\ (c)}$ + Ingestion $volume_{(a)}$]. Multiple sets of data were compared using single-factor ANOVA coupled with the Scheffe's post hoc test. n=12. Statistical significance compared with the controls is indicated by the asterisks. Means ± SEMs. $^{**}p < 0.01$.

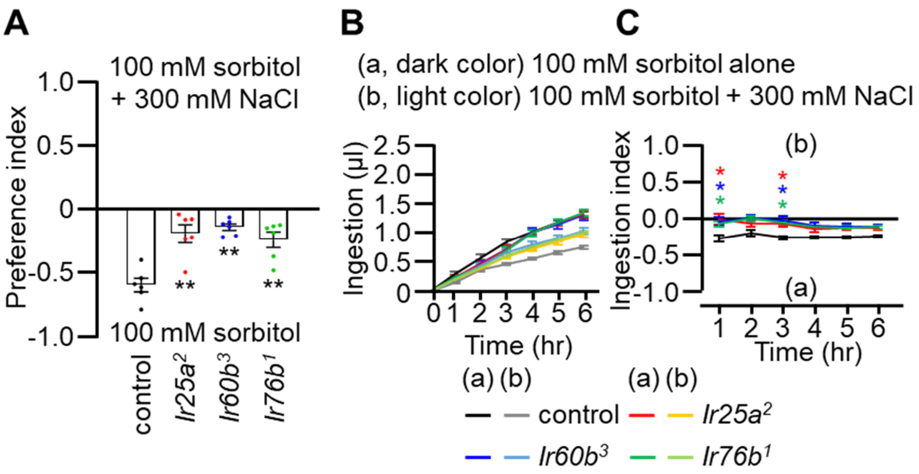


**Figure 3—figure supplement 2.** Two-way solid-food choice assay and DrosoX binary capillary feeding assay using 100 mM sorbitol with or without 300 mM NaCl. (**A**) Two-way solid-food choice assay to assess whether the *Ir25a²*, *Ir60b³*, and *Ir76b¹* mutants exhibit a deficit in high salt avoidance. The flies were given a choice between 100 mM sorbitol versus 100 mM sorbitol plus 300 mM NaCl in alternating wells of microtiter dishes. n=6. (**B** and **C**) DrosoX assays used to test the relative volumes consumed by control, *Ir25a²*, *Ir60b³*, and *Ir76b¹* flies when presented with capillaries containing either 100 mM sorbitol (a) or 100 mM sorbitol plus 300 mM NaCl (b). n=12. (**B**) Volumes of each of two food options. (**C**) Ingestion indexes (I.I) to indicate the relative consumption of the two foods. I.I formula: [Ingestion volume$_{(b)}$ – Ingestion volume$_{(a)}$]/ [Ingestion volume$_{(b)}$ + Ingestion volume$_{(a)}$]. Multiple sets of data were compared using single-factor ANOVA coupled with Scheffe's post hoc test. Statistical significance compared with the controls. Means ± SEMs. *$p < 0.05$. **$p < 0.01$.

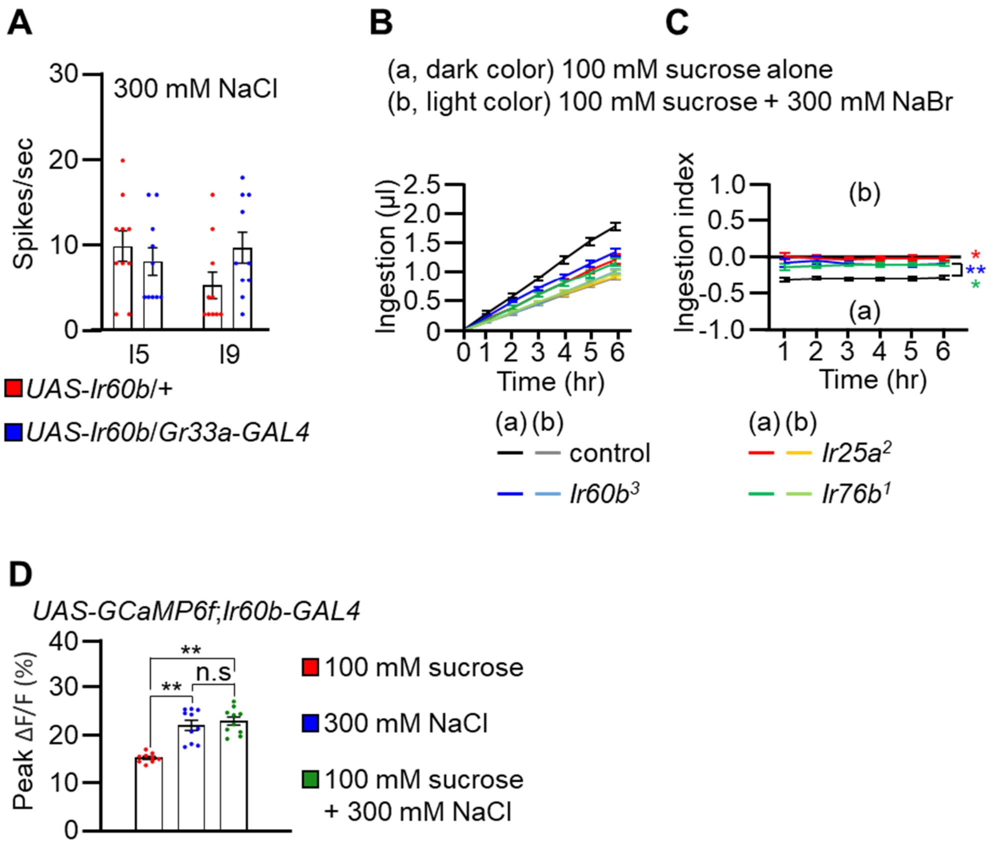


**Figure 4—figure supplement 1.** Testing whether ectopic expression of *Ir60b* confers responses to 300 mM NaCl, measurement of intake of sucrose plus 300 mM NaBr using the DrosoX assay, and $Ca^{2+}$ response of *Ir60b* GRNs. (**A**) Tip recordings in response to 300 mM NaCl were performed on I5 and I9 sensilla of the indicated flies. *UAS-Ir60b* was expressed in Class B GRNs under control of the *Gr33a-GAL4*. n=10. (**B** and **C**) Droso-X assays used to test the relative volumes consumed by *Ir25a²*, *Ir60b³*, and *Ir76b¹* flies when presented with capillaries containing either 100 mM sucrose (a) or 100 mM sucrose plus 300 mM NaBr (b). n=12. (**B**) Volumes of each of two food options. (**C**) Ingestion indexes (I.I) to indicate the relative consumption of the two foods. I.I formula: [Ingestion volume$_{(b)}$ – Ingestion volume$_{(a)}$]/ [Ingestion volume$_{(b)}$ + Ingestion volume$_{(a)}$]. (**D**) GCaMP6f responses of *Ir60b* GRNs to sucrose only, NaCl only, or a combination of sucrose and NaCl. n=10. Multiple sets of data were compared using single-factor ANOVA coupled with Scheffe's post hoc test. Statistical significance compared with the controls. Means ± SEMs. *$p < 0.05$. **$p < 0.01$.

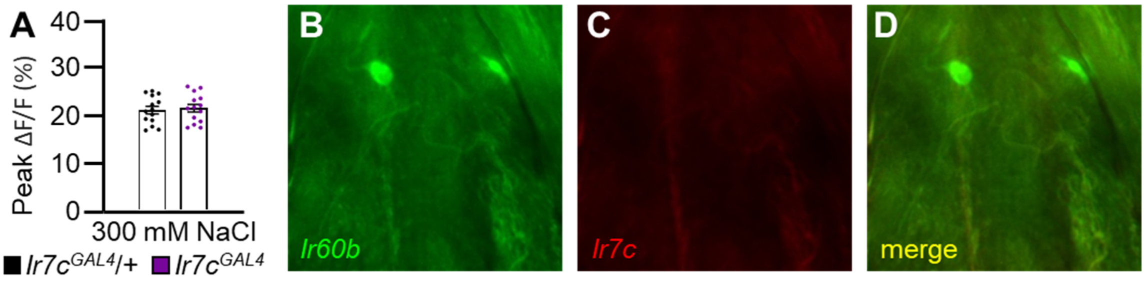


**Figure 4—figure supplement 2.** GCaMP6f responses evoked by 300 mM NaCl in *Ir60b* GRNs of control and $Ir7c^{GAL4}$ flies, and relative expression of *Ir60b* and *Ir7c* reporters in the LSO. (**A**) GCaMP6f responses were stimulated by 300 mM NaCl in *Ir60b* GRNs from the $Ir7c^{GAL4}$ mutant ($Ir7c^{GAL4}$;*UAS-GCaMP6f*/+;*Ir60b-GAL4*/+) and from the $Ir7c^{GAL4}$/+ control ($Ir7c^{GAL4}$/+;*UAS-GCaMP6f*/+;*Ir60b-GAL4*/+). n=8–10. The pairwise comparison was conducted using an unpaired Student's *t*-test. Means ± SEMs. (**B—D**) Testing for expression of the *Ir7c* reporter in *Ir60b* GRNs. The *Ir7c* reporter consisted of a gene encoding GAL4 fused to RFP, and expressed under the direct control of the *Ir7c* promoter ($Ir7c^{GAL4::VP166\text{-}RFP}$)[1]. The *Ir60b* reporter consisted of *UAS-GFP* driven by the *Ir60b-GAL4*. The genotype of the flies was $Ir7c^{GAL4::VP166\text{-}RFP}$;*Ir60b-GAL4*;*UAS-mCD8*::*GFP* . The *Ir60b* reporter was detected with anti-GFP, and the *Ir7c* reporter was detected with anti-DsRed.
